# Retention and data exclusion challenges for representative longitudinal neuroimaging in the understanding of addiction

**DOI:** 10.1101/2025.09.04.674267

**Authors:** Jocelyn A. Ricard, Russell A. Poldrack, Keith Humphreys

**Author notes:** **Correspondence:** Jocelyn A. Ricard, Stanford University, Stanford Neurosciences Interdepartmental Program, Stanford University School of Medicine, 290 Jane Stanford Way, Stanford, CA 94305.

## Abstract

Head motion during resting-state functional magnetic resonance imaging (rsfMRI) poses a major challenge for neuroimaging research, often leading to data quality concerns and participant exclusions, particularly among pediatric and clinical populations. Although necessary for ensuring reliable data, motion-related exclusions may inadvertently bias samples by disproportionately excluding certain sociodemographic groups. Using data from the Adolescent Brain Cognitive Development (ABCD) Study, we employed both frequentist and Bayesian approaches to examine how head motion-related exclusions and participant retention shape sample composition over time. At baseline (ages 9-10), Black, Hispanic, and Asian youth were significantly more likely than White youth to be excluded due to excessive head motion; these disparities were not observed at the two-year follow-up (ages 11-13). In contrast, disparities in retention persisted; Black participants were less likely to return for follow-up, even after accounting for socioeconomic factors and motion. Together, these findings highlight how both motion-related exclusions and differential retention can systematically influence the representativeness of longitudinal neuroimaging samples, with important implications for the generalizability of research in addiction neuroscience.

## Introduction

Scientists support the fundamental principle that the benefits of human subjects research should be shared by all (cf. the principle of Justice outlined in the Belmont Report) (U.S. Department of Health and Human Services, 1979). This is especially relevant in studying addiction, because the spread of synthetic opioids like fentanyl, compounded by the COVID pandemic, has widened racial disparities in overdose mortality in the United States (Han et al., 2022). Understanding the nature of and best treatments for addiction for all Americans is made more challenging by the exclusion criteria for many research studies (e.g., residential instability) disproportionately excluding Black, low-income, and/or individuals with more severe cases of substance use disorder (SUD) (Humphreys, 2000; Humphreys et al., 2007).

The exclusion of participants for excessive head motion in MRI studies is an example of a research practice that may disproportionately exclude certain sociodemographic groups, causing the final analytic sample to diverge demographically from the initially recruited one (Dhamala et al., 2025; Ricard et al., 2023). Although some exclusion criteria are necessary for safety (e.g., metal as an exclusion criterion in an MRI study), overly broad exclusion criteria that are not undergirded by safety concerns and disproportionately exclude vulnerable populations have been common in addiction research (e.g., clinical trials (Humphreys, 2000)). Here we examine the ways in which head motion exclusions may have a similar effect in MRI research, unintentionally perpetuating disparities that may endanger the validity and generalizability of findings across different populations. If head motion exclusion used in MRI studies disproportionately exclude vulnerable individuals, who are often the most affected by these disparities, it may compromise the generalizability of models relating brain to behavior in high-motion populations (A. S. Greene et al., 2022; D. J. Greene et al., 2016; Ramduny et al., 2025).

Head motion correction is a critical preprocessing step for ensuring good data quality in neuroimaging research, particularly for resting-state functional connectivity (Dosenbach et al., 2017; Satterthwaite et al., 2012; Van Dijk et al., 2012). Exclusion of individuals with high head motion can reduce confounding effects related to motion artifacts and ensure good data quality by enhancing signal-to-noise ratio, thereby improving validity of the resulting inferences. As scientists, we, of course, understand the necessity of addressing this issue. However, the tradeoff is that prior work has established relationships between head motion and both sociodemographic and psychological variables in MRI studies, suggesting that when we exclude individuals through data quality steps, we disproportionately exclude both demographically diverse individuals and patients (Cosgrove et al., 2022) as well as introducing problematic selection biases.

In line with the Belmont principles, major efforts have been made to establish large-population human biomedical datasets. One example of this is the Adolescent Brain Cognitive Development Study (ABCD Study), which is the largest study assessing brain development in the United States (Casey et al., 2018). Given a key initial goal of the ABCD Study’s sampling strategy was to create a population-relevant cohort in order to examine the developmental pathways of substance use (Compton et al., 2019; Karcher & Barch, 2021), this dataset provides a unique opportunity to investigate these questions. Recent comprehensive work has documented this issue at baseline in the ABCD Study(Cosgrove et al., 2022; Peverill et al., 2025), demonstrating that high-motion status is systematically related to a wide array of sociodemographic and clinical variables. However, a critical unanswered question is how sociodemographic factors and head motion patterns are associated with exclusion and participant retention over the course of ABCD’s follow-up study participation. Therefore, in this present study, we first examined how the first-level quality control inclusion criterion for resting-state functional magnetic resonance imaging (rsfMRI) and subsequent high-head motion exclusions for subjects who pass first-level quality control relate to sociodemographics at baseline. We then examine how exclusion based on high-motion status is associated with exclusion and participant retention over the ABCD Study’s 2-year follow-up. This longitudinal approach allows us to assess the dynamic relationships between motion-related exclusion, study retention, sample representativeness, and generalizability.

## Methods

### Participants

Data were obtained from the ABCD Study Data Release 5.1, a large, longitudinal, population-based cohort in the United States (Casey et al., 2018). Our analyses focused on participants with available rsfMRI data at baseline (ages 9-10) and the 2-year follow-up (ages 11-13). The ABCD Data Analysis, Informatics, and Resource Center (DAIRC) provides a first-level quality control variable for resting fMRI data, based on a comprehensive quality control (QC) pipeline. We first analyzed all subjects (N=10,280 [1,588 removed to missing data]) to assess the relationship between this QC-based exclusion criterion and presence within the 2-year follow-up dataset; although missing data in the follow-up timepoint dataset does not necessarily imply study loss to follow-up (e.g. it could simply reflect follow-up that was too late to be included in the 5.1 data release), for the purposes of this analysis, we assume that non-retention causes are random and thus attribute missingness to failed retention. We then generated our primary sample using methods similar to Cosgrove et al., 2022, We then subsetted the data to include only those with complete data (i.e., participants with complete data across all sociodemographic and head motion variables of interest) at baseline, which resulted in a baseline sample of N = 10,280. We further included all participants with a recommendation for rsfMRI inclusion by the ABCD DAIRC (imgincl_rsfmri_include = 1) (Hagler Jr et al., 2019; Power et al., 2012), which left us with a final baseline sample of N = 8561. A CONSORT flow diagram detailing baseline, follow-up, and longitudinal sample participant numbers is presented in **Fig. s1**.

We also generated a follow-up sample, and a longitudinal sample for direct comparison between the baseline and 2-year follow-up timepoints, using only participants who had available data at both timepoints. The DAIRC inclusion criteria and subsetting to those with complete data at the 2-year follow-up for the relevant variables resulted in a final 2-year follow-up sample size of N = 5789. A longitudinal sample was created using only participants who had available data at both the baseline and 2-year follow-up timepoints. We performed listwise deletion, retaining only those participants with complete data across all sociodemographic and head motion variables of interest. This resulted in a complete longitudinal sample of N = 4,658. Participant race and ethnicity were parent-reported at baseline and categorized as White, Black, Hispanic, Asian, and Other, consistent with ABCD Study conventions.

### High-Head Motion and Exclusion Status

To identify participants with excessive in-scanner head motion for resting-state scans, we categorized high-motion individuals as those whose mean framewise displacement (FD) across all sessions was ≥ 0.2 mm. This threshold aligns with prior subject-level exclusion using the ABCD Study (Marek et al., 2019), and is consistent with the ABCD Study’s processing pipeline, which excludes time points with FD ≥ 0.2 mm from resting-state functional connectivity analyses to reduce residual motion artifacts (Hagler Jr et al., 2019; Power et al., 2014). Individuals with a mean FD ≥ 0.2 mm were defined as the high-motion or ‘excluded’ group for our analyses, whereas those with a mean FD < 0.2 mm were defined as the low-motion or ‘included’ group.

The analyses in the present report were also conducted using samples defined by different thresholds (mean FD ≥ 0.15 mm and ≥ 0.3 mm); the results were largely similar to those reported at the FD ≥ 0.2 mm threshold (see **Figs. s1** and **s2**).

### Analyses

All analyses were conducted in R (version 4.4.2); the full analysis code is available at (https://github.com/ricardjocelyn/head-motion-exclusion; doi: 10.5281/zenodo.17049858). First, to meet the Data Analysis, Informatics, & Resource Center (DAIRC) inclusion criterion, rsfMRI and corresponding T1w scans must pass raw and manual QC, retain at least 375 frames after motion censoring, have valid unwarp and registration metrics, meet coverage thresholds, and produce non-missing DAIRC-derived correlation maps. Given that this initial quality control criterion already excludes some individuals on the basis of high-motion, and researchers often impose additional, more stringent motion exclusion criteria on top of this, we wanted to assess whether our additional step introduced a selection bias into the final sample. Next, we investigated sociodemographic predictors of both data exclusion and sample retention using a series of generalized linear mixed-effects models (GLMERs) with the lme4 package (Bates et al., 2015). To account for possible clustered data structure, the longitudinal model included random intercepts for study site and for subject nested within family ID, and the baseline and follow-up models included random intercepts for study site and family ID, except when those led to failures of model convergence. Covariates were standardized in order to improve model convergence.

Because large samples like ABCD can result in highly significant statistical results even when the effect size is small, we also quantified evidence for/against the null using Bayes factors. These were computed using the BayesTestR package (Makowski et al., 2019), which uses an approximation to the Bayes Factor based on the Bayesian Information Criterion (BIC) (Wagenmakers, 2007). Bayes factors are reported as BF_10_; values greater than one reflect evidence in favor of the alternative hypothesis (values around 1 considered anecdotal, 3–10 moderate, 10–30 strong, and >30 very strong evidence), while values less than one reflect evidence in favor of the null (Kass & Raftery, 1995). Effect sizes for mixed-effect logistic regression were quantified using the pseudo R-squared computed using the r.squaredGLMM function from the MuMin R package (Barton & Barton, 2015).

## Results

### Analysis of racial/ethnic disparities in the DAIRC inclusion criteria

First, we fit a generalized linear mixed-effects model predicting the binary DAIRC inclusion variable (imgincl_rsfmri_include) to assess whether the initial quality control criteria had a racially disparate effect (N = 10280). The model included fixed effects for household income, parental education, sex, age, and race/ethnicity, with random intercepts for family ID and site to account for nested data structure.

Black participants were significantly less likely to be included compared to White participants based upon the DAIRC-criterion, but the Bayes factor supported the null (OR = 0.79, 95% CI [0.66, 0.95], p = 0.012, BF_10_ = 0.21). However, a likelihood ratio test comparing the full model to a reduced model without the race/ethnicity term was not statistically significant (χ^2^(4) = 8.76, *p* = .07, BF_10_ = 7.54e-07), with the Bayes factor providing strong evidence that the race/ethnicity variable did not improve model fit. Given that the DAIRC exclusion criteria do not have a racially divergent impact, the subsequent analyses used the DAIRC-included sample as used by Cosgrove et al., 2022. Full and reduced model parameters are provided in the **Supplementary Tables s1 and s2**.

### Head motion and sociodemographics as independent correlates of retention

We first examined the association between head motion and retention, adjusting for socioeconomic and demographic covariates. Starting with the most complex model, we tested the interaction between head motion and race, controlling for income, parental education, gender, and age (Supplemental Table S5). The interaction was not statistically significant (χ^2^(4)

= 6.55, p = .16), and none of the individual interaction terms (of mean motion with each race category) had a significant p-value. Further, the Bayes factor strongly supported excluding the interaction (BF_10_ = 3.61e-07).

Next, we assessed the role of race/ethnicity in predicting follow-up participation using both frequentist and Bayesian approaches. Compared to White participants, the odds of retention were significantly lower for Black individuals (OR = 0.63, 95% CI [0.52, 0.75], p < .001) and for those in the Other race category (OR = 0.80, 95% CI [0.67, 0.95], p = .012), whereas no other race groups differed significantly from White participants. These effects are illustrated in **Fig. 1** in terms of predicted probabilities (probability of retention), adjusted for all continuous covariates at their mean values for race/ethnicity and mean head motion.

**Figure 1.**
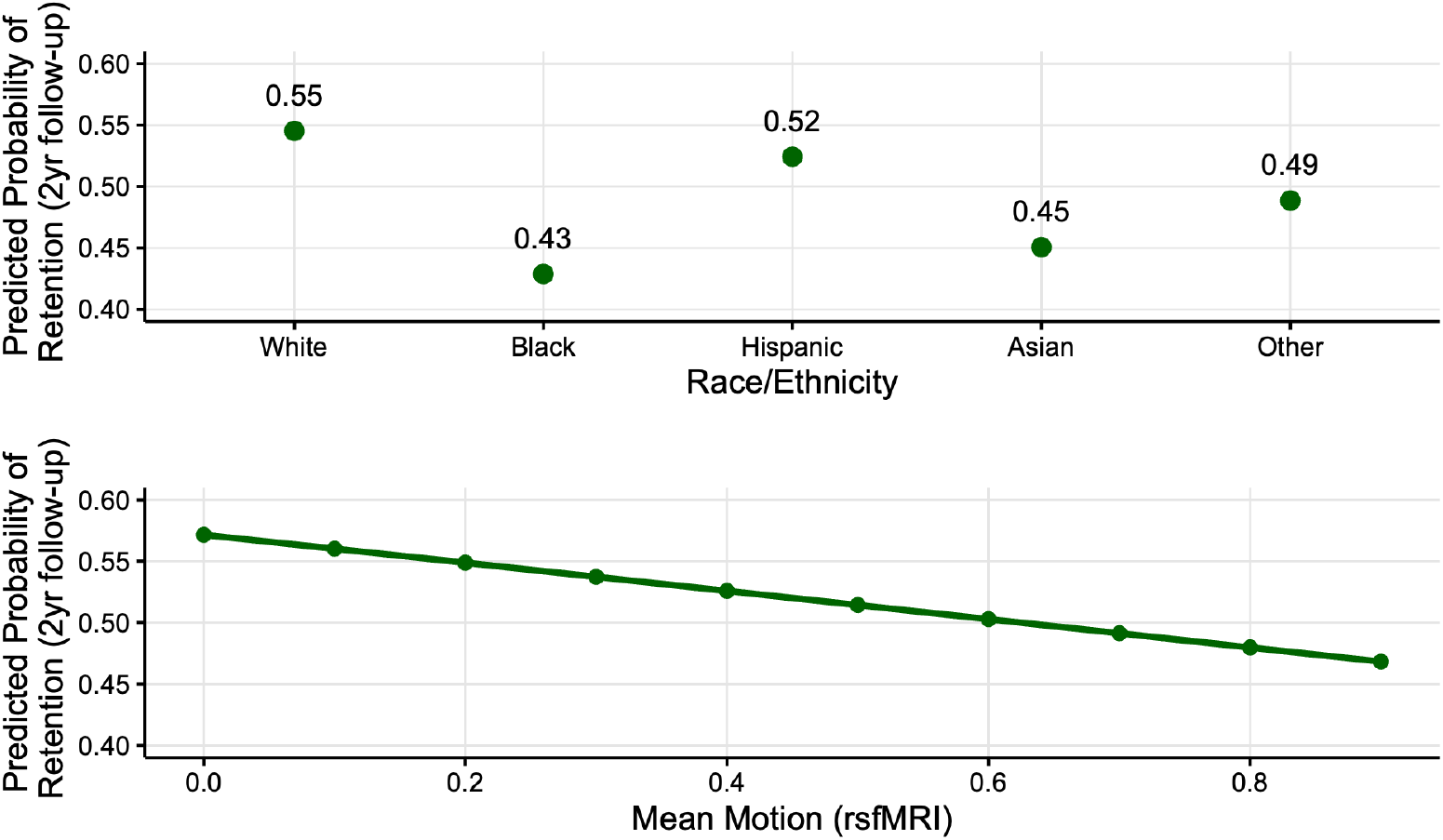
Predicted probabilities of having follow-up data across race/ethnicity and mean rsfMRI motion at baseline.

To further evaluate the importance of individual race categories, we calculated Bayes factors comparing models that combined each specific race group with White versus the full model that treated all race categories separately. The Bayes factor for Black was 2189.6, indicating strong evidence that Black participants should be modeled separately from White participants. In contrast, Hispanic, Asian, and Other groups showed BF_10_ values of 0.0177, 0.068, and 0.256 respectively, indicating little evidence to model these categories separately.

A likelihood ratio test comparing the full model (including all race indicators) to a reduced model (excluding race) also supported the inclusion of race overall (χ^2^(4) = 29.6, p = < 0.001). However, a BIC-based Bayes factor comparing these two models yielded BF_10_ ≈ 0.04 in favor of the reduced model, suggesting that from a Bayesian model selection perspective, including all race indicators may not be justified by the increase in model complexity. The parameter estimates for other covariates were similar in the reduced model (**Table 1**, right side).

**Table 1.** Odds ratios, 95% CIs and p-values for the full and reduced retention models. The raw parameter estimates are supplied in the Supplemental Tables 1 and 2.

|  | Full model |  |  |  | Reduced Model |  |  |  |
| --- | --- | --- | --- | --- | --- | --- | --- | --- |
| predictor | OR | 95% CI | p-value | $BF_{10}$ | OR | 95% CI | p-value | $BF_{10}$ |
| income_z | 1.092 | (1.015, 1.175) | 0.018 | 0.176 | 1.155 | (1.077, 1.239) | 0.000 | 38.22 |
| parent_ed_z | 1.069 | (0.993, 1.150) | 0.076 | 0.051 | 1.073 | (0.999, 1.154) | 0.054 | 0.068 |
| rsfmri_meanmotion | 0.631 | (0.496, 0.802) | 0.000 | 12.66 | 0.618 | (0.486, 0.785) | 0.000 | 25.36 |
| race/ethnicity Black | 0.626 | (0.519, 0.754) | 0.000 | 2189.6 |  |  |  |  |
| race/ethnicity Hispanic | 0.919 | (0.778, 1.085) | 0.319 | 0.0177 |  |  |  |  |
| race/ethnicity Asian | 0.684 | (0.464, 1.007) | 0.054 | 0.068 |  |  |  |  |
| race/ethnicity Other | 0.797 | (0.668, 0.950) | 0.012 | 0.256 |  |  |  |  |
| female | 0.819 | (0.739, 0.907) | 0.000 | 16.82 | 0.814 | (0.735, 0.901) | 0.000 | 26.55 |
| age_z | 0.944 | (0.897, 0.994) | 0.028 | 0.119 | 0.943 | (0.895, 0.992) | 0.023 | 0.138 |

**Table 2.** Odds ratios, 95% CIs and p-values for the <u>baseline</u> high-motion status exclusion full and reduced models. The parameter estimates are supplied in the Supplemental Tables 4 and 5.

| predictor | Full model |  |  |  | Reduced Model |  |  |  |
| --- | --- | --- | --- | --- | --- | --- | --- | --- |
| | OR | 95% CI | p-value | $BF_{10}$ | OR | 95% CI | p-value | $BF_{10}$ |
| income_z | 0.927 | [0.869, 0.989] | 0.023 | 0.144 | 0.901 | [0.847, 0.959] | 0.001 | 2.48 |
| parent_ed_z | 0.903 | [0.846, 0.965] | 0.002 | 1.09 | 0.885 | [0.830, 0.943] | 0.000 | 12.06 |
| race/ethnicity Black | 1.231 | [1.044, 1.451] | 0.013 | 0.232 |  |  |  |  |
| race/ethnicity Hispanic | 1.342 | [1.157, 1.557] | 0.000 | 20.819 |  |  |  |  |
| race/ethnicity Asian | 1.503 | [1.066, 2.118] | 0.020 | 0.156 |  |  |  |  |
| race/ethnicity Other | 1.031 | [0.879, 1.210] | 0.707 | 0.012 |  |  |  |  |
| female | 0.736 | [0.671, 0.808] | 0.000 | 1.67e+07 | 0.739 | [0.673, 0.810] | 0.000 | 1.1e+07 |
| age_z | 0.783 | [0.746, 0.821] | 0.000 | 6.39e+20 | 0.783 | [0.747, 0.821] | 0.000 | 6.59e+20 |

Taken together, these results illustrate the complexity of modeling race/ethnicity. Although the overall contribution of race to model fit is ambiguous across inferential frameworks, there is consistent and strong evidence that Black participants are substantially less likely to have follow-up data compared to white participants, even after adjusting for socioeconomic and demographic factors.

### Racial and ethnic disparities in motion-based data exclusion are present at baseline, but not at the 2-year follow-up

Next, we assessed the role of race/ethnicity in predicting exclusion due to high-motion status at baseline using both frequentist and Bayesian approaches. At baseline, exclusion due to high-motion rates on the full sample (n = 8561) were highest for Black participants (48.3%), followed by Hispanic (44.8%), Asian (43.5%), Other (37.9%) and White (35.4%) participants (**Fig. 2**, top). At the 2-year follow-up (n = 5789), overall exclusion rates declined across all groups, consistent with decreased motion across child development as well as increased exclusion and loss to followup for high-motion individuals. Rates remained highest among Black (33.5%) and Asian (29.7%) participants, followed by Other (28.1%), Hispanic (26.3%), and White (24%) participants (**Fig. 2**, bottom). Demographic breakdowns for the full sample at baseline and follow-up are described in **Table 5** and **Table 6**, respectively.

**Figure 2.**
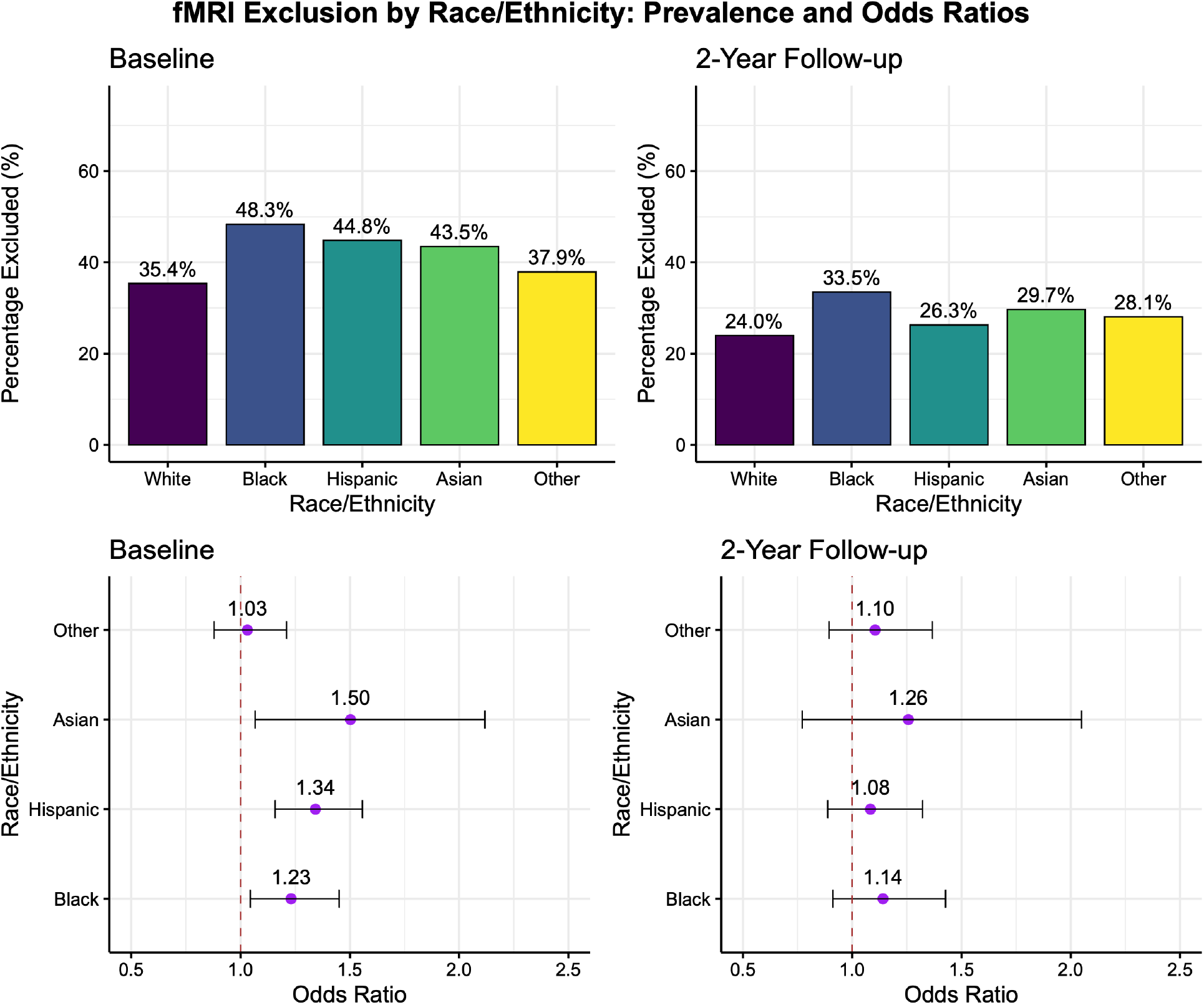
Bar graphs represent the percentage of participants excluded from analyses due to high head motion at baseline and 2-year follow-up, calculated as the proportion of excluded participants within each racial/ethnic group. Forest plots show odds ratios with 95% confidence intervals relative to the White participant reference group. Data were obtained from Adolescent Brain and Cognitive Development Study (ABCD), with exclusion thresholds set at a mean rsfMRI framewise displacement (FD) ≥ 0.2 mm. Racial/ethnic groups are labeled according to the ABCD Study categories.

To identify predictors of data exclusion due to high head motion in the baseline data, we fitted a series of generalized linear mixed-effects models modeling exclusion based on high-motion status as a function of racial/ethnic group with covariates including household income, parental education, age, and sex while also accounting for random intercepts for study site and family ID. At baseline (ages 9-10), there were significant racial/ethnic disparities in exclusion based on high-motion status. Consistent with prior work examining high-head motion status (Cosgrove et al., 2022), a model comparison showed that including race/ethnicity significantly outperformed the model without race/ethnicity (χ^2^(4) = 21.13, p < .001); here again the Bayes.

Factor preferred the reduced model (BF_10_ = 5.2e-04), suggesting that the significant effect nonetheless provides weak evidence for the alternative hypothesis. Group-specific estimates revealed that, relative to White participants, the odds of being excluded for high head motion were significantly higher for Black (OR = 1.23, 95% CI [1.04, 1.45], p = .013), Hispanic (OR = 1.34, 95% CI [1.16, 1.56], p < .001), and Asian (OR = 1.50, 95% CI [1.07, 2.12], p = .02) participants. This was not the case for participants who identified as ‘Other’ (OR = 1.03, CI [0.88, 1.21], p = 0.71). These effects are illustrated in **Fig. 3** in terms of predicted probabilities (probability of exclusion), adjusted for all continuous covariates at their mean values for race/ethnicity and mean head motion.

**Fig. 3.**
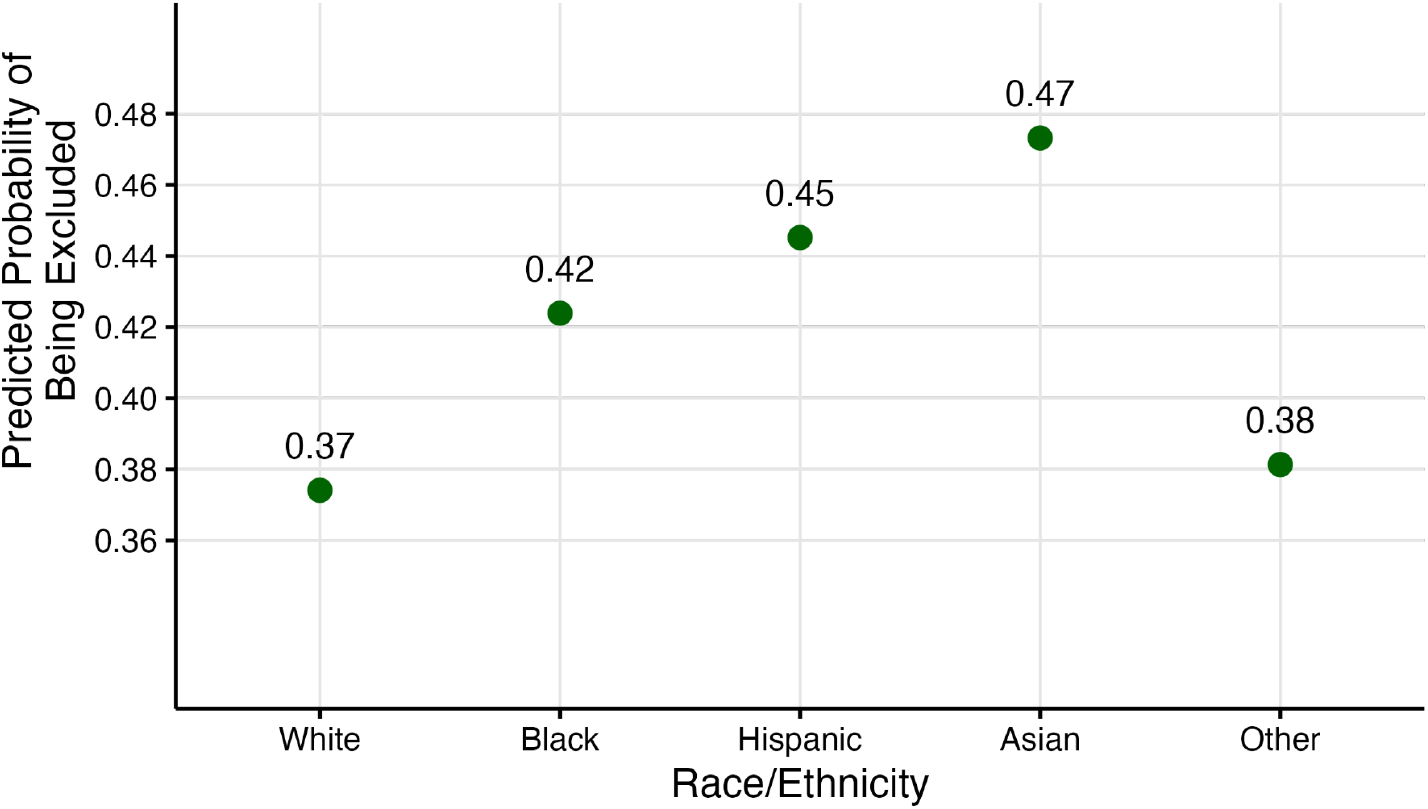
Predicted probabilities of being excluded due to high-motion status at baseline across race/ethnicity.

In the 2-year follow-up data (ages 11-13), these racial/ethnic disparities in high-head motion status exclusion were no longer present. The GLMER analysis at the 2-year follow-up revealed no significant differences in the odds of exclusion based on high-head motion status for any racial/ethnic group compared to White participants (χ^2^(4) = 2.5, p = 0.64, BF_10_ = 1.04e-07, in favor of the reduced model).

A longitudinal model examining youth with data at both timepoints (n = 4658) simultaneously found that after controlling for covariates (sex, age, parental education, household income), race/ethnicity was not a statistically significant predictor of high-motion status. A model comparison confirmed that adding race/ethnicity did not significantly improve model fit (χ^2^(4) = 5.81, p = 0.214, BF_10_ = 2.10e-07 in favor of the reduced model), suggesting no overall effect of race/ethnicity across the two timepoints (**Table 4**).

**Table 3.** Odds ratios, 95% CIs and p-values for the <u>follow-up</u> high-motion status exclusion full and reduced models. The parameter estimates are supplied in the Supplemental Tables 6 and 7.

|  | Full model |  |  |  | Reduced Model |  |  |  |
| --- | --- | --- | --- | --- | --- | --- | --- | --- |
| predictor | OR | 95% CI | p-value | BF<sub>10</sub> | OR | 95% CI | p-value | BF<sub>10</sub> |
| income_z | 0.937 | [0.861, 1.020] | 0.134 | 0.04 | 0.921 | [0.850, 0.998] | 0.045 | 0.096 |
| parent_ed_z | 0.955 | [0.877, 1.041] | 0.296 | 0.023 | 0.951 | [0.874, 1.035] | 0.244 | 0.026 |
| race/ethnicity Black | 1.141 | [0.912, 1.427] | 0.249 | 0.0254 |  |  |  |  |
| race/ethnicity Hispanic | 1.083 | [0.888, 1.322] | 0.429 | 0.1794 |  |  |  |  |
| race/ethnicity Asian | 1.257 | [0.771, 2.049] | 0.359 | 0.0197 |  |  |  |  |
| race/ethnicity Other | 1.105 | [0.894, 1.366] | 0.357 | 0.0199 |  |  |  |  |
| female | 0.568 | [0.500, 0.645] | < 0.001 | 1.13e+15 | 0.569 | [0.501, 0.647] | < 0.001 | 8.46e+14 |
| age_z | 0.814 | [0.763, 0.868] | < 0.001 | 5.22e+06 | 0.814 | [0.763, 0.869] | < 0.001 | 4.58e+06 |

**Table 4.** Odds ratios, 95% CIs and p-values for the <u>longitudinal</u> high-motion status exclusion full and reduced models. The parameter estimates are supplied in the Supplemental Tables 8 and 9.

|  | Full model |  |  |  | Reduced Model |  |  |  |
| --- | --- | --- | --- | --- | --- | --- | --- | --- |
| predictor | OR | 95% CI | p-value | BF<sub>10</sub> | OR | 95% CI | p-value | BF<sub>10</sub> |
| income_z | 0.965 | [0.882, 1.055] | 0.431 | 0.014 | 0.937 | [0.860, 1.020] | 0.133 | 0.032 |
| parent_ed_z | 0.918 | [0.838, 1.006] | 0.066 | 0.056 | 0.902 | [0.825, 0.987] | 0.025 | 0.129 |
| race/ethnicity Black | 1.262 | [0.992, 1.607] | 0.059 | 0.078 |  |  |  |  |
| race/ethnicity Hispanic | 1.221 | [0.995, 1.499] | 0.056 | 0.067 |  |  |  |  |
| race/ethnicity Asian | 1.159 | [0.680, 1.976] | 0.588 | 0.013 |  |  |  |  |
| race/ethnicity Other | 1.057 | [0.847, 1.320] | 0.622 | 0.013 |  |  |  |  |
| female | 0.588 | [0.517, 0.669] | < 0.001 | 4.68e+12 | 0.590 | [0.519, 0.671] | < 0.001 | 3.0e+12 |
| study visit baseline | 2.698 | [2.412, 3.018] | < 0.001 | 1.39e+72 | 2.700 | [2.413, 3.020] | < 0.001 | 1.81e+72 |
| age_z | 0.763 | [0.715, 0.814] | < 0.001 | 1.26e+13 | 0.764 | [0.716, 0.815] | < 0.001 | 9.88e+12 |

**Table 5.**
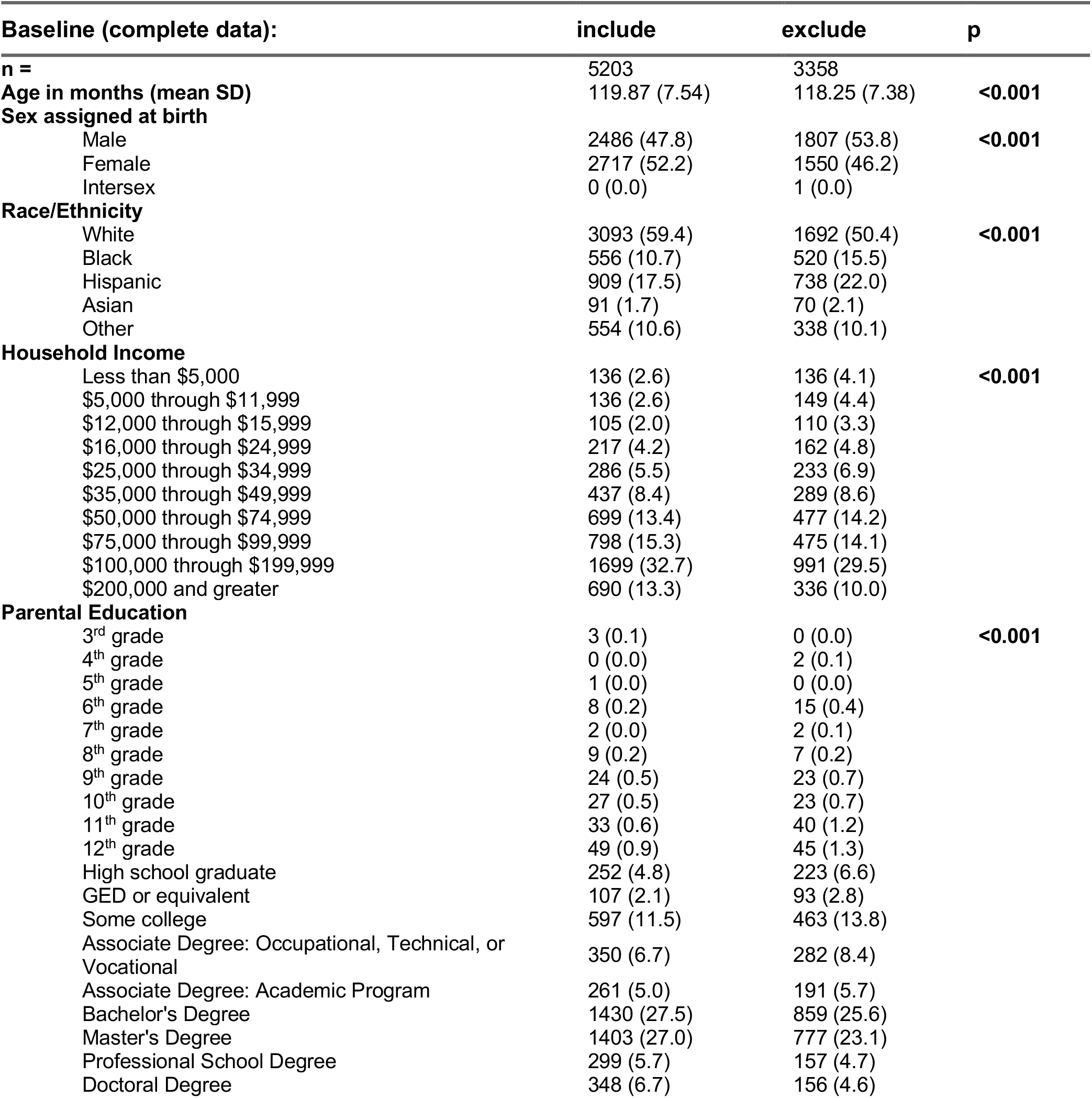
Demographic characteristics of the included (n=5203) and excluded (n=3358) sample in the ABCD Study at baseline at mean FD ≥ 0.2 mm threshold.

**Table 6.** Demographic characteristics of the included (n=4287) and excluded (n=1502) sample in the ABCD Study at year-2 follow-up at mean FD ≥ 0.2 mm threshold.

| **2-year Follow Up (complete data):** | **include** | **exclude** | **p** |
| --- | --- | --- | --- |
| **n =** | 4287 | 1502 |  |
| **Age in months (mean SD)** | 143.78 (7.86) | 142.30 (7.41) | **<0.001** |
| **Sex assigned at birth** |  |  |  |
| Male | 2140 (49.9) | 930 (61.9) | **<0.001** |
| Female | 2147 (50.1) | 572 (38.1) |  |
| **Race/Ethnicity** |  |  |  |
| White | 2581 (60.2) | 813 (54.1) | **<0.001** |
| Black | 427 (10.0) | 215 (14.3) |  |
| Hispanic | 795 (18.5) | 283 (18.8) |  |
| Asian | 64 (1.5) | 27 (1.8) |  |
| Other | 420 (9.8) | 164 (10.9) |  |
| **Household Income** |  |  |  |
| Less than \$5,000 | 80 (1.9) | 60 (4.0) | **<0.001** |
| \$5,000 through \$11,999 | 106 (2.5) | 46 (3.1) | |
| \$12,000 through \$15,999 | 69 (1.6) | 31 (2.1) | |
| \$16,000 through \$24,999 | 147 (3.4) | 52 (3.5) | |
| \$25,000 through \$34,999 | 231 (5.4) | 89 (5.9) | |
| \$35,000 through \$49,999 | 287 (6.7) | 124 (8.3) | |
| \$50,000 through \$74,999 | 599 (14.0) | 205 (13.6) | |
| \$75,000 through \$99,999 | 638 (14.9) | 203 (13.5) | |
| \$100,000 through \$199,999 | 1510 (35.2) | 508 (33.8) | |
| \$200,000 and greater | 620 (14.5) | 184 (12.3) | |
| **Parental Education** |  |  |  |
| 4<sup>th</sup> grade | 1 (0.0) | 0 (0.0) | **0.003** |
| 6<sup>th</sup> grade | 6 (0.1) | 2 (0.1) |  |
| 7<sup>th</sup> grade | 3 (0.1) | 1 (0.1) |  |
| 8<sup>th</sup> grade | 9 (0.2) | 0 (0.0) |  |
| 9<sup>th</sup> grade | 26 (0.6) | 8 (0.5) |  |
| 10<sup>th</sup> grade | 25 (0.6) | 11 (0.7) |  |
| 11<sup>th</sup> grade | 28 (0.7) | 17 (1.1) |  |
| 12<sup>th</sup> grade | 42 (1.0) | 13 (0.9) |  |
| High school graduate | 197 (4.6) | 96 (6.4) |  |
| GED or equivalent | 75 (1.7) | 41 (2.7) |  |
| Some college | 516 (12.0) | 211 (14.0) |  |
| Associate degree: Occupational, Technical, or Vocational | 308 (7.2) | 113 (7.5) |  |
| Associate degree: Academic Program | 240 (5.6) | 85 (5.7) |  |
| Bachelor's Degree | 1196 (27.9) | 386 (25.7) |  |
| Master's Degree | 1124 (26.2) | 389 (25.9) |  |
| Professional School Degree | 250 (5.8) | 58 (3.9) |  |
| Doctoral Degree | 241 (5.6) | 71 (4.7) |  |

However, there was a significant effect of study visit on high-motion status, with participants more likely to be excluded at the baseline visit compared to the two-year follow-up (β = 0.99, z = 17.35, p < .001, BF_10_ = 1.39e+72 favoring the full model), consistent with a reduction in head motion across development and greater failure to retain individuals with higher trait levels of head motion.

## Discussion

The present study combined cross-sectional and longitudinal analyses to examine relationships between head motion exclusion and demographics in the ABCD study. We replicated prior findings (Cosgrove et al., 2022; Peverill et al., 2025) by demonstrating that minoritized youth were more likely to exceed motion-based exclusion criteria at baseline. However, we find that these data-quality disparities were not present at the 2-year follow-up. This difference stands in contrast to the persistent disparities in overall study retention, where Black and female participants remained significantly more likely to drop out at the 2-year follow-up, even after accounting for individual differences in baseline head motion. This key distinction suggests that the mechanisms contributing to poor in-scanner data quality, which may be ameliorated with age and experience (D. J. Greene et al., 2016), may be distinct from the more entrenched, systemic factors driving long-term study disengagement. Our analyses reveal the particular vulnerability of Black participants who were not only more likely to have high head motion and be flagged for exclusion at baseline, but also were more likely to not return for the 2-year follow-up study visit. The results also highlight the complex nature of data missingness within large datasets like ABCD Study over time.

Populations with neuropsychiatric diagnoses tend to exhibit significantly more in-scanner head motion than healthy control subjects (D. J. Greene et al., 2016; Makowski et al., 2019). Given that one of the primary goals of the ABCD study was to investigate the impact of substance use and child development, this potential bias has concrete implications. Imagine a researcher has recruited participants for a study examining the impact of substance use on cognitive function compromising a racially/ethnically diverse group, and employs high head motion as an exclusion criterion. Prior to head motion exclusion, the results show nuanced differences in cognitive function across racial/ethnic groups and between patients and healthy controls. However, because of higher head motion observed predominantly among Black individuals, and also more broadly among clinical populations (Andre et al., 2015; Makowski et al., 2019), the effective sample composition changes after exclusions, leading to a reduction in the diversity of the sample. If, for instance, the initial analysis showed that in Black patients, substance use-related cognitive function declines were closely related to specific patterns of substance use, but this relationship was not observed in White patients, the study’s conclusions might be skewed by the exclusions. After excluding participants with high head motion, a criterion that predominantly affects the overall sample composition, leading to an increased representation of White, educated, and socioeconomically advantaged individuals, the study might incorrectly conclude that the observed decline in cognitive function is universally unrelated to specific patterns of substance use, reducing the generalizability of estimates in the group.

This scenario emphasizes the risk of systematic bias, hence not fully representing the spectrum of effects across different racial/ethnic groups. The study’s conclusions may not accurately reflect how substance use-induced cognitive impairments manifest in populations facing SUDs. The present work thus underscores the critical need for inclusive research methodologies that accurately reflect the diverse populations affected by SUDs.

How should we address this problem? First, it’s essential that any associations between MRI signals and clinical or behavioral variables are tested in diverse populations and replicated across independent samples to ensure that observed effects are not confined to a specific demographic group or experimental setting (preferably with pre-registered sampling and analysis plans). Second, studies should report whether observed associations are robust to different motion correction strategies, and report the demographics and behavioral features of those subjects who are excluded due to head motion and/or lost to follow-up. An example of this approach in the context of structural MRI has recently shown how associations between brain structure and clinical features can differ substantially under different exclusion criteria (Elyounssi et al., 2025), highlighting the importance of assessing these differences across analysis workflows. Any associations that are highly sensitive to preprocessing choices should be viewed with caution, and any substantive conclusions drawn from them should remind readers that the findings are not universal but limited to the non-excluded groups (Humphreys & Williams, 2018). Third, in studies of group differences it is essential to characterize how exclusion criteria change the group membership. For example, in the context of autism it has been shown that motion exclusion results in a systematically different autism population compared to the full sample. Any inferences based on such a sample are liable to the well-known “matching fallacy” (Meehl, 1970).

MRI has played a pivotal role in our understanding of SUDs in the brain. For example, MRI studies have pinpointed prefrontal cortex dysfunction to be associated with greater likelihood of relapse and elucidate predictors of recovery outcomes for individuals, e.g., (Goldstein & Volkow, 2011; Yip et al., 2019). But, by definition, MRI findings can only be assumed to apply to the people who provide the data for MRI research. Because decisions about whom to enroll and exclude in research inherently involve tradeoffs, explicitly acknowledging such tradeoffs and their implications should be a standard research practice. Concretely, this means that if the head motion criteria and adjustments employed in a study result in the under-representation of individuals of racial and ethnic backgrounds, this limitation should be discussed in the paper. Many groups have attempted to highlight and create guidelines to deal with the best practices in inclusion-exclusion of those who participate in our MRI studies (D. J. Greene et al., 2016; Li et al., 2022; Ramduny et al., 2025; Satterthwaite et al., 2013). Although motion distortion and artifact correction are unavoidable, these goals can come in conflict when researchers aim to incorporate groups that are disproportionately left out due to exclusion criteria in neuroimaging studies.

Our analyses also highlight another challenge of working with large datasets like ABCD: even very small effects can be highly statistically significant cf. (Dick et al., 2021). In this study we used multiple approaches to triangulate the assessment of associations between our variables of interest: null hypothesis statistical testing, effect size estimates to quantify the magnitude of the effect (both relative, as in odds ratios, and absolute, as in r-squared measures), and model comparisons using Bayes factors. Together these highlighted the high statistical power of ABCD, and also the fact that methods with different assumptions may not agree, as when frequentist and Bayesian analyses differed in their conclusions Echoing Dick et al. (2021), we think that a more nuanced approach to statistical hypothesis testing is necessary in the context of large datasets like ABCD.

### Limitations

Several limitations should be considered when interpreting these findings. First, our longitudinal models were necessarily restricted to participants with complete data at both timepoints, introducing selection bias. Our own results demonstrate that participants who dropped out were systematically different, exhibiting higher baseline head motion and being more likely to identify as Black. The resulting range restriction may explain the lack of effects in the longitudinal sample. Consequently, the resolution of data quality disparities we observed in our retained sample may not fully generalize to the more vulnerable populations that are most likely to leave longitudinal studies over time. Second, it is crucial to remember that race and ethnicity are social constructs that reflect a complex interplay of unmeasured sociocultural, environmental, and systemic factors that shape a child’s experience and behavior in a research setting, and the observed disparities should be interpreted in this light (Bryant et al., 2022). Furthermore, the scope of this study was limited to the transition into early adolescence (ages 9-13). The predictors of head motion and their relationship with retention may evolve as new challenges, such as the emergence of psychiatric illness, e.g., substance use become more salient in mid-to-late adolescence (Bachman et al., 2018; Zapert et al., 2002). The re-emergence or transformation of these disparities in later years remains an open question for future research.

Lastly, the 2-year follow-up data are not complete, and missingness at the 2-year follow-up does not necessarily reflect lack of retention. For example, data at the 2-year follow-up may have been missing because youth missed visits, withdrew from the study, or had late visits that were not captured at the time of this release (Ewing et al., 2022). Despite these limitations, this study provides critical, novel evidence that sources of sampling bias in neuroimaging are not static. It highlights the necessity for researchers to move beyond cross-sectional snapshots of data quality and instead longitudinally assess and report how exclusion procedures and retention shape sample composition over time, thereby ensuring the generalizability and equity of their findings.

## Acknowledgments

This work was supported by: Stanford University Knight-Hennessy Scholars Program (JAR); National Academies of Sciences, Engineering, and Medicine’s Ford Foundation Predoctoral Fellowship (JAR).

