## Supplementary Materials for "Retention and data exclusion challenges for representative longitudinal neuroimaging in the understanding of addiction"

**Correspondence:**

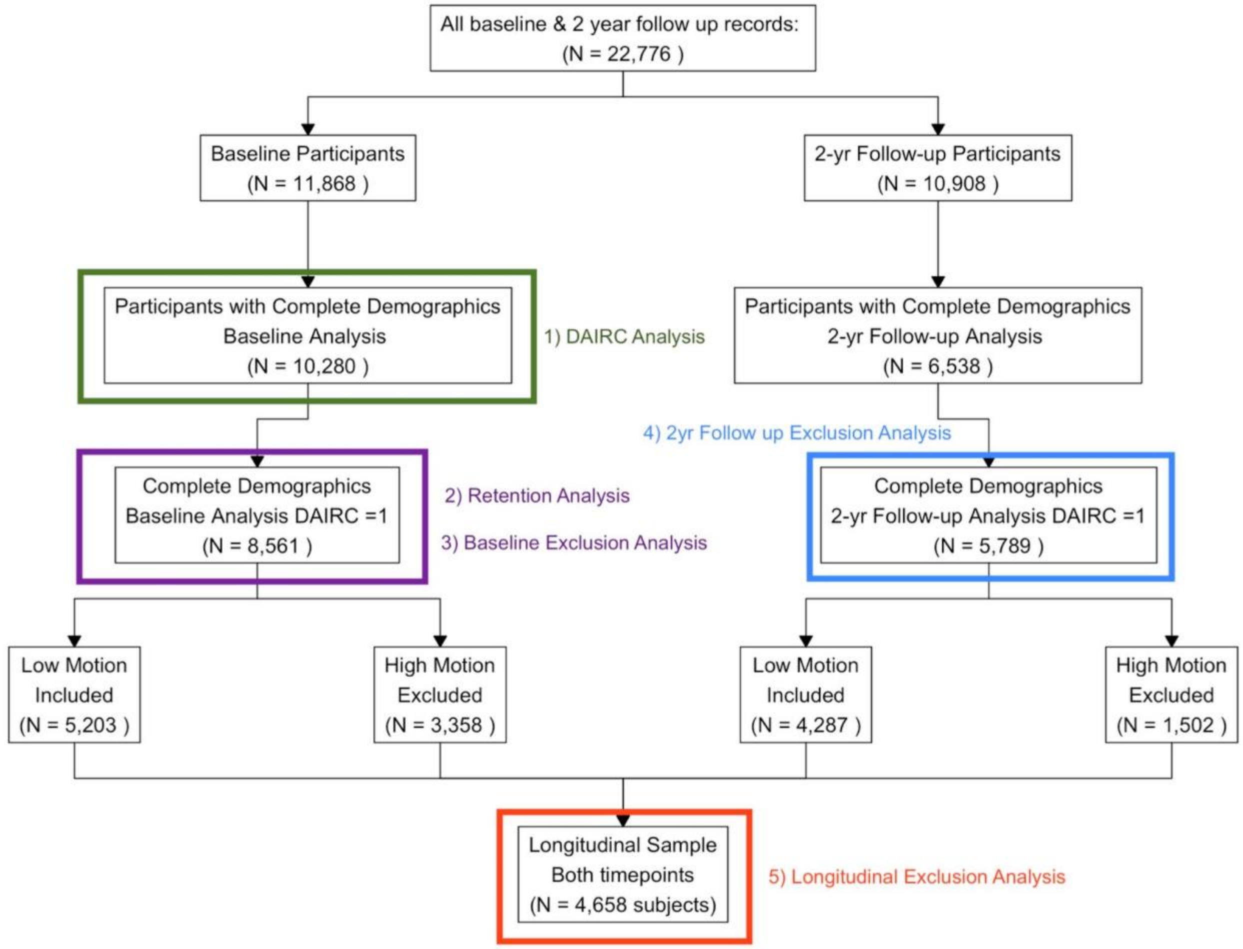


**Figure s1:** CONSORT flow diagram detailing baseline (N = 8561 participants), follow-up (N = 5789 participants), and longitudinal (N = 4658 participants) samples. The longitudinal sample only includes participants with complete data at both baseline and 2-year follow-up timepoints. High-motion was classified as mean FD $\geq$ 0.2 mm. Low-motion was classified as mean FD < 0.2 mm.

**DAIRC Analysis:**

Full DAIRC model:

| predictor | estimate | estimate 95% CI | OR | OR 95% CI | Std. error | p-value | $BF_{10}$ |
|---|---|---|---|---|---|---|---|
| income_z | 0.139 | (0.063, 0.215) | 1.149 | (1.065, 1.24) | 0.039 | <0.001 | 4.64 |
| parent_ed_z | 0.066 | (-0.012, 0.143) | 1.068 | (0.988, 1.154) | 0.040 | 0.096 | 0.038 |
| female | 0.530 | (0.418, 0.642) | 1.699 | (1.519, 1.901) | 0.057 | <0.001 | 7.20e+16 |
| race/ethnicity Black | -0.232 | (-0.413, -0.051) | 0.793 | (0.662, 0.95) | 0.092 | 0.012 | 0.209 |
| race/ethnicity Hispanic | 0.028 | (-0.152, 0.208) | 1.029 | (0.859, 1.232) | 0.092 | 0.758 | 0.010 |
| race/ethnicity Asian | -0.175 | (-0.562, 0.212) | 0.839 | (0.57, 1.236) | 0.197 | 0.375 | 0.143 |
| race/ethnicity Other | 0.022 | (-0.168, 0.212) | 1.022 | (0.845, 1.236) | 0.097 | 0.821 | 0.010 |
| age_z | 0.207 | (0.151, 0.264) | 1.231 | (1.163, 1.302) | 0.029 | <0.001 | 1.52e+09 |

**Table S1. Full model:** glmer(imgincl_rsfmri_include ~ income_z + parent_ed_z + female + race_ethnicity_label + age_z + (1 | rel_family_id) + (1 | site). The random intercept variance was 0.180 for rel_family_id and 0.376 for site.

Reduced DAIRC model:

| predictor | estimate | estimate 95% CI | OR | OR 95% CI | Std. error | p-value | $BF_{10}$ |
|---|---|---|---|---|---|---|---|
| income_z | 0.166 | (0.093, 0.239) | 1.181 | (1.098, 1.27) | 0.037 | <0.001 | 159.3 |
| parent_ed_z | 0.064 | (-0.012, 0.14) | 1.066 | (0.988, 1.15) | 0.039 | 0.098 | 0.037 |
| female | 0.527 | (0.415, 0.639) | 1.694 | (1.515, 1.894) | 0.057 | <0.001 | 4.6e+16 |
| age_z | 0.207 | (0.15, 0.263) | 1.229 | (1.162, 1.301) | 0.029 | <0.001 | 1.27e+09 |

**Table S2**. **Reduced model:** glmer(imgincl_rsfmri_include ~ income_z + parent_ed_z + female + age_z + (1 | rel_family_id) + (1 | site). The random intercept variance was 0.179 for rel_family_id and 0.383 for site.

### Retention:

| predictor | estimate | 95% CI | std.error | p-value |
|---|---|---|---|---|
| income_z | 0.144 | [0.074, 0.214] | 0.037 | 0.000 |
| parent_ed_z | 0.071 | [-0.001, 0.143] | 0.038 | 0.054 |
| rsfmri_meanmotion | -0.482 | [-0.722, -0.242] | 0.123 | 0.000 |
| female | -0.206 | [-0.308, -0.104] | 0.052 | 0.000 |
| age_z | -0.059 | [-0.110, -0.008] | 0.026 | 0.023 |

**Table S3.** Fixed effects estimates from the repeated measures logistic regression reduced model: glmer(has_followup_data ~ income_z + parent_ed_z + rsfmri_meanmotion + female + age_z + (1 | rel_family_id) + (1 | site)). The random intercept variance was 0.672 for rel_family_id and 0.295 for site.

| predictor | estimate | 95% CI | std.error | p-value |
|---|---|---|---|---|
| income_z | 0.088 | [0.015, 0.162] | 0.037 | 0.018 |
| parent_ed_z | 0.067 | [-0.007, 0.140] | 0.038 | 0.076 |
| rsfmri_meanmotion | -0.461 | [-0.702, -0.221] | 0.123 | 0.000 |
| race/ethnicity Black | -0.469 | [-0.655, -0.282] | 0.095 | 0.000 |
| race/ethnicity Hispanic | -0.085 | [-0.251, 0.082] | 0.085 | 0.319 |
| race/ethnicity Asian | -0.380 | [-0.768, 0.007] | 0.198 | 0.054 |
| race/ethnicity Other | -0.228 | [-0.404, -0.051] | 0.090 | 0.012 |
| female | -0.200 | [-0.302, -0.098] | 0.052 | 0.000 |
| age_z | -0.058 | [-0.109, -0.006] | 0.026 | 0.028 |

**Table S4.** Fixed effects estimates from the repeated measures logistic regression full model: glmer(has_followup_data ~ income_z + parent_ed_z + rsfmri_meanmotion + race_ethnicity_label + female + age_z + (1 | rel_family_id) + (1 | site)). The random intercept variance was 0.673 for rel_family_id and 0.284 for site.

| predictor | estimate | 95% CI | std.error | p-value |
|---|---|---|---|---|
| income_z | 0.087 | [0.014, 0.160] | 0.037 | 0.020 |
| parent_ed_z | 0.068 | [-0.005, 0.142] | 0.038 | 0.068 |
| rsfmri_meanmotion | -0.339 | [-0.685, 0.006] | 0.176 | 0.054 |
| race/ethnicity Black | -0.463 | [-0.716, -0.210] | 0.129 | 0.000 |
| race/ethnicity Hispanic | 0.006 | [-0.217, 0.229] | 0.114 | 0.960 |
| race/ethnicity Asian | 0.193 | [-0.443, 0.830] | 0.325 | 0.552 |
| race/ethnicity Other | -0.224 | [-0.471, 0.023] | 0.126 | 0.076 |
| female | -0.200 | [-0.302, -0.098] | 0.052 | 0.000 |
| age_z | -0.057 | [-0.108, -0.006] | 0.026 | 0.029 |
| rsfmri_meanmotion:Black | -0.045 | [-0.692, 0.602] | 0.330 | 0.891 |
| rsfmri_meanmotion:Hispanic | -0.372 | [-0.992, 0.248] | 0.316 | 0.239 |
| rsfmri_meanmotion:Asian | -2.620 | [-5.009, -0.230] | 1.219 | 0.032 |
| rsfmri_meanmotion:Other | -0.025 | [-0.777, 0.728] | 0.384 | 0.949 |

**Table S5.** Fixed effects estimates from the repeated measures logistic regression model with an interaction term: glmer(has_followup_data ~ income_z + parent_ed_z + rsfmri_meanmotion * race_ethnicity_label + female + age_z + (1 | rel_family_id) + (1 | site). The random intercept variance was 0.672 for rel_family_id and 0.285 for site.

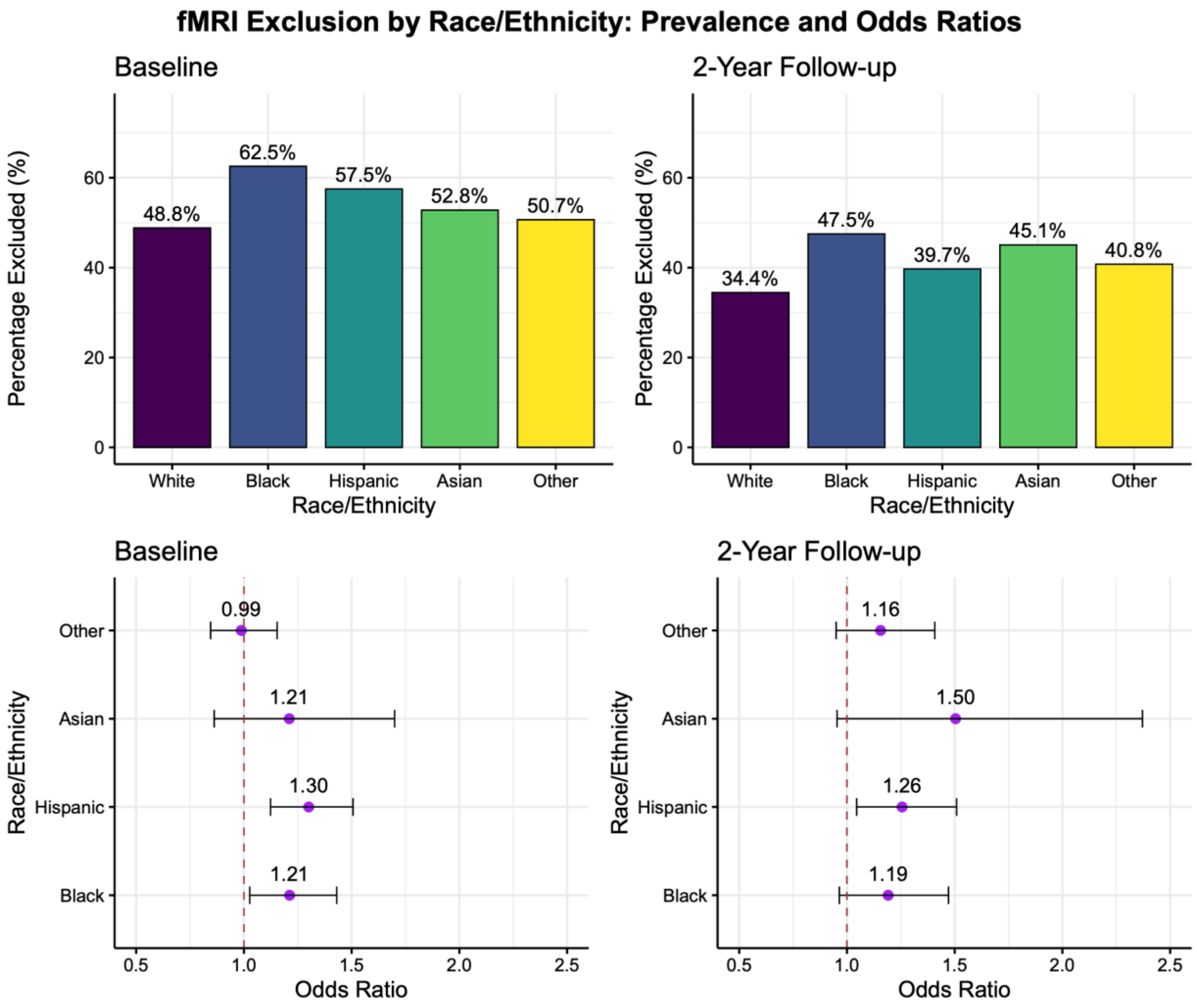


**Figure s2:** Bar graphs represent the percentage of participants excluded from analyses due to high head motion at baseline and 2-year follow-up, calculated as the proportion of excluded participants within each racial/ethnic group. Forest plots show odds ratios with 95% confidence intervals relative to the White participant reference group. Data were obtained from Adolescent Brain and Cognitive Development Study (ABCD), with exclusion thresholds set at a mean rsfMRI framewise displacement *(FD)* $\geq$ *0.15 mm*. Racial/ethnic groups are labeled according to the ABCD Study categories.

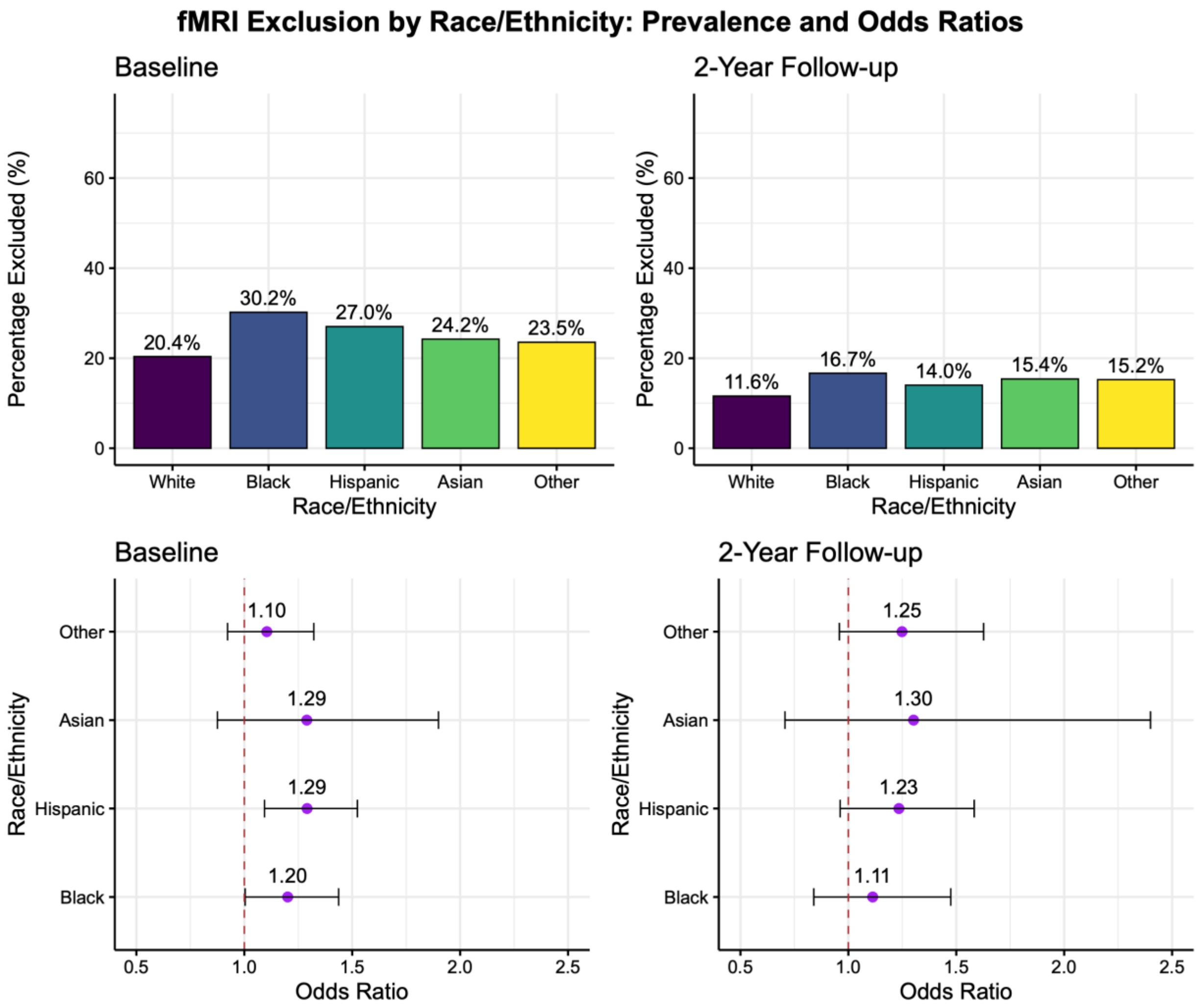


**Figure s3:** Bar graphs represent the percentage of participants excluded from analyses due to high head motion at baseline and 2-year follow-up, calculated as the proportion of excluded participants within each racial/ethnic group. Forest plots show odds ratios with 95% confidence intervals relative to the White participant reference group. Data were obtained from Adolescent Brain and Cognitive Development Study (ABCD), with exclusion thresholds set at a mean rsfMRI framewise displacement *(FD)* $\geq$ *0.3 mm*. Racial/ethnic groups are labeled according to the ABCD Study categories.

**Exclusion:**

Baseline models:

| predictor | estimate | 95% CI | std.error | p-value |
|---|---|---|---|---|
| income_z | -0.10425 | [-0.166, -0.042] | 0.03159 | 0.001 |
| parent_ed_z | -0.12255 | [-0.187, -0.059] | 0.03274 | < 0.000 |
| female | -0.30284 | [-0.395, -0.210] | 0.0472 | < 0.000 |
| age_z | -0.24457 | [-0.292, -0.197] | 0.0243 | < 0.000 |

**Table S6.** Fixed effects estimates from the repeated measures baseline logistic regression reduced model: glmer(exclude ~ income_z + parent_ed_z + female + age_z + (1 | rel_family_id) + (1 | site)). The random intercept variance was 0.132 for rel_family_id and 0.102 for site.

| predictor | estimate | 95% CI | std.error | p-value |
|---|---|---|---|---|
| income_z | -0.07571 | [-0.141, -0.011] | 0.03321 | 0.02264 |
| parent_ed_z | -0.10152 | [-0.167, -0.036] | 0.03343 | 0.00239 |
| race/ethnicity Black | 0.20781 | [0.043, 0.372] | 0.08386 | 0.01321 |
| race/ethnicity Hispanic | 0.29448 | [0.146, 0.443] | 0.07565 | < 0.000 |
| race/ethnicity Asian | 0.40730 | [0.064, 0.750] | 0.17502 | 0.01996 |
| race/ethnicity Other | 0.03075 | [-0.129, 0.191] | 0.08168 | 0.70656 |
| female | -0.30626 | [-0.399, -0.214] | 0.04727 | < 0.000 |
| age_z | -0.24480 | [-0.292, -0.197] | 0.02432 | < 0.000 |

**Table S7.** Fixed effects estimates from the repeated measures baseline logistic regression full model: glmer(exclude ~ income_z + parent_ed_z + race_ethnicity_label + female + age_z + (1 | rel_family_id) + (1 | site)). The random intercept variance was 0.131 for rel_family_id and 0.104 for site.

Follow-up models:

| predictor | estimate | 95% CI | std.error | p-value |
|---|---|---|---|---|
| income_z | -0.082 | [-0.162, -0.002] | 0.04087 | 0.045 |
| parent_ed_z | -0.050 | [-0.135, 0.034] | 0.04307 | 0.244 |
| female | -0.563 | [-0.135, 0.034] | 0.06496 | < 0.000 |
| age_z | -0.205 | [-0.270, -0.141] | 0.03298 | < 0.000 |

**Table S8.** Fixed effects estimates from the repeated measures logistic follow-up regression reduced model: glmer(exclude ~ income_z + parent_ed_z + female + age_z + (1 | rel_family_id) + (1 | site)). The random intercept variance was 0.127 for rel_family_id and 0.187 for site.

| predictor | estimate | 95% CI | std.error | p-value |
|---|---|---|---|---|
| income_z | -0.06486 | [-0.149, 0.020] | 0.04324 | 0.134 |
| parent_ed_z | -0.04586 | [-0.132, 0.040] | 0.04384 | 0.296 |
| race/ethnicity Black | 0.13175 | [-0.092, 0.356] | 0.11422 | 0.249 |
| race/ethnicity Hispanic | 0.08014 | [-0.118, 0.279] | 0.10139 | 0.429 |
| race/ethnicity Asian | 0.22876 | [-0.260, 0.717] | 0.24930 | 0.359 |
| race/ethnicity Other | 0.09981 | [-0.112, 0.312] | 0.10829 | 0.357 |
| female | -0.56579 | [-0.693, -0.438] | 0.06502 | < 0.000 |
| age_z | -0.20614 | [-0.271, -0.141 | 0.03299 | < 0.000 |

**Table S9.** Fixed effects estimates from the repeated measures follow-up logistic regression full model: glmer(exclude ~ income_z + parent_ed_z + race_ethnicity_label + female + age_z + (1 | rel_family_id) + (1 | site)). The random intercept variance was 0.128 for rel_family_id and 0.183 for site.

Longitudinal models:

| predictor | estimate | 95% CI | std.error | p-value |
|---|---|---|---|---|
| income_z | -0.06552 | [-0.151, 0.020] | 0.04364 | 0.1332 |
| parent_ed_z | -0.10283 | [-0.192, 0.014] | 0.04579 | 0.0247 |
| female | -0.52771 | [-0.656, -0.399] | 0.06563 | < 0.000 |
| study visit baseline | 0.99315 | [0.881, 1.105] | 0.05719 | < 0.000 |
| age_z | -0.26935 | [-0.334, -0.205] | 0.03304 | < 0.000 |

**Table S10.** Fixed effects estimates from the repeated measures longitudinal logistic regression reduced model: glmer(exclude ~ income_z + parent_ed_z + female + eventname + age_z + (1 | rel_family_id/src_subject_id) + (1 | site)). The random intercept variance was 0.767 for rel_family_id, 0.156 for site, and 0.539 for src_subject_id:rel_family_id.

| predictor | estimate | 95% CI | std.error | p-value |
|---|---|---|---|---|
| income_z | -0.03610 | [-0.335, -0.205] | 0.04588 | 0.4313 |
| parent_ed_z | -0.08580 | [-0.177, 0.005] | 0.04665 | 0.0659 |
| race/ethnicity Black | 0.23278 | [-0.009, 0.474] | 0.12312 | 0.0587 |
| race/ethnicity Hispanic | 0.19959 | [-0.005, 0.404] | 0.10456 | 0.0563 |
| race/ethnicity Asian | 0.14759 | [-0.386, 0.681] | 0.27213 | 0.5876 |
| race/ethnicity Other | 0.05582 | [-0.166, 0.278] | 0.11334 | 0.6223 |
| female | 0.04588 | [-0.083, 0.175] | 0.06565 | < 0.000 |
| study visit baseline | 0.99237 | [0.880, 1.104] | 0.05719 | < 0.000 |
| age_z | -0.27025 | [-0.335, -0.205] | 0.03304 | < 0.000 |

**Table S11.** Fixed effects estimates from the repeated measures longitudinal logistic regression full model: glmer(exclude ~ income_z + parent_ed_z + female + race_ethnicity_label + eventname + age_z + (1 | rel_family_id/src_subject_id) + (1 | site)). The random intercept variance was 0.7631 for rel_family_id, 0.1509 for site, and 0.5401 for src_subject_id:rel_family_id.
